# tDCS induced GABA change is associated with the simulated electric field in M1, an effect mediated by grey matter volume in the MRS voxel

**DOI:** 10.1101/2022.04.27.489665

**Authors:** Tulika Nandi, Oula Puonti, William T. Clarke, Caroline Nettekoven, Helen C. Barron, James Kolasinski, Taylor Hanayik, Emily L. Hinson, Adam Berrington, Velicia Bachtiar, Ainslie Johnstone, Anderson M. Winkler, Axel Thielscher, Heidi Johansen-Berg, Charlotte J. Stagg

## Abstract

**Background and Objective:** Transcranial direct current stimulation (tDCS) has wide ranging applications in neuro-behavioural and physiological research, and in neurological rehabilitation. However, it is currently limited by substantial inter-subject variability in responses, which may be explained, at least in part, by anatomical differences that lead to variability in the electric field (E-field) induced in the cortex. Here, we tested whether the variability in the E-field in the stimulated cortex during tDCS, estimated using computational simulations, explains the variability in tDCS induced changes in GABA, a neurophysiological marker of stimulation effect.

**Methods:** Data from five previously conducted MRS studies were combined. The anode was placed over the left primary motor cortex (M1, 3 studies, N = 24) or right temporal cortex (2 studies, N = 32), with the cathode over the contralateral supraorbital ridge. Single voxel spectroscopy was performed in a 2×2×2cm voxel under the anode in all cases. MRS data were acquired before and either during or after 1mA tDCS using either a sLASER sequence (7T) or a MEGA-PRESS sequence (3T). sLASER MRS data were analysed using LCModel, and MEGA-PRESS using FID-A and Gannet. E-fields were simulated in a finite element model of the head, based on individual MPRAGE images, using SimNIBS. Separate linear mixed effects models were run for each E-field variable (mean and 95th percentile; magnitude, and components normal and tangential to grey matter surface, within the MRS voxel). The model included effects of time (pre or post tDCS), E-field, grey matter volume in the MRS voxel, and a 3-way interaction between time, E-field and grey matter volume. Additionally, we ran a permutation analysis using PALM to determine whether E-field anywhere in the brain, not just in the MRS voxel, correlated with GABA change.

**Results:** In M1, higher mean E-field magnitude was associated with greater tDCS-induced decreases in GABA (t(24) = 3.24, p = 0.003). Further, the association between mean E-field magnitude and GABA change was moderated by the grey matter volume in the MRS voxel (t(24) = −3.55, p =0.002). These relationships were consistent across all E-field variables except the mean of the normal component. No significant relationship was found between tDCS-induced GABA decrease and E-field in the temporal voxel. No significant clusters were found in the whole brain analysis.

**Conclusions:** Our data suggest that the electric field induced by tDCS within the brain is variable, and is significantly related to tDCS-induced decrease in GABA, a key neurophysiological marker of stimulation. These findings strongly support individualised dosing of tDCS, at least in M1. Further studies examining E-fields in relation to other outcome measures, including behaviour, will help determine the optimal E-fields required for any desired effects.

**Highlights:**

- We study the link between individually simulated electric field dose and tDCS-induced change in GABA in the cortex.
- The electric field strength in the brain correlates with a decrease in GABA in the motor cortex.
- The correlation between the electric field and GABA change is modulated by the amount of grey matter in the MRS voxel.
- We find no association between the electric field and GABA in the temporal cortex.

## INTRODUCTION

Transcranial direct current stimulation (tDCS) shows promise as a potential therapeutic intervention for a range of neurological and psychiatric conditions (Indahlastari et al., 2021; Kang et al., 2016). However, the current evidence for clinical application of tDCS is deemed to be ineffective or only probably effective (Lefaucheur et al., 2017). One factor limiting clinical translation of tDCS is the high inter-subject variability in response (Chew et al., 2015; López-Alonso et al., 2015; Wiethoff et al., 2014). Such variability may be caused by a variety of factors, including trait differences in anatomy and neurophysiology between subjects, or the prevailing brain state during tDCS application (Woods et al., 2016). However, while the brain state may be experimentally controlled or accounted for, anatomical differences between subjects cannot be reduced. It has therefore been suggested that individual anatomy should be accounted for when dosing tDCS (Evans et al., 2020).

The effect of individual anatomy on the electric field (E-field) distribution in the brain can be studied using simulations (Huang et al., 2019; Saturnino, Puonti, et al., 2019; Thielscher et al., 2015) that rely on realistic volume conductor models of the head anatomy, which are constructed from a structural MRI scan of a subject – a ‘head model’ (Nielsen et al., 2018; Puonti, Van Leemput, et al.,2020). With an appropriate head model, the electric field induced in the brain by different tDCS electrode configurations can be modelled using the finite element method (FEM) (Saturnino, Madsen, et al., 2019). Simulations conducted on multiple subjects have shown that anatomical differences such as the thickness of the CSF layer, scalp to coil distance, and local cortical folding all influence the E-field induced in the underlying cortex (Laakso et al., 2015; Opitz et al., 2015; Puonti, Saturnino, et al., 2020; Thielscher et al., 2011). While the need for individually-tailored stimulation protocols is widely recognized (Hartwigsen et al., 2015; Krause & Cohen Kadosh, 2014; Liu et al., 2018), most tDCS studies still apply the same extracranial current amplitude for all subjects leading to a large range of E-field magnitudes in the cortex (Antonenko et al., 2019; Mezger et al., 2021). Several approaches have been suggested to reduce the E-field variability across subjects, with the implicit assumption that this also leads to a reduced variability in *responses*, ranging from tuning the current amplitude (Caulfield et al., 2020) to an optimization of both the electrode locations and input currents (Dmochowski et al., 2011; Saturnino, Siebner, et al., 2019). However, it remains uncertain whether the E-field is a significant predictor of neurophysiological outcomes of interest (Antonenko et al.,2019).

Several studies have demonstrated that anodal tDCS decreases GABA in the stimulated cortex (Bachtiar et al., 2015, 2018; Barron et al., 2016; Koolschijn et al., 2019; Stagg et al., 2011), a proposed mechanism through which tDCS promotes plasticity and learning (Kolasinski et al., 2019). Indeed, the magnitude of tDCS-induced decreases in GABA predict behaviour (Kim et al., 2014; O’Shea et al., 2014; Stagg et al.,2011). MRS-assessed GABA might therefore act as an individual marker of behaviourally-relevant, neurophysiological effects of tDCS.

In this paper, we combined data from several previously conducted tDCS-MRS studies, to test the hypothesis that inter-individual differences in E-field are correlated with the tDCS-induced GABA decrease, such that greater E-fields would lead to greater tDCS-induced GABA decreases. We examined the E-field components normal and tangential to the grey matter surface, in addition to the E-field magnitude, since in vitro data suggest that the E-field direction relative to the neuronal axis has a large impact on tDCS effects (Radman et al., 2009). We concentrated on the primary motor cortex (M1), as the majority of previous studies investigating tDCS induced GABA change have focussed on this region. To determine whether any relationships demonstrated in M1 were also found in other cortical regions, we also included data from studies that used an MRS voxel in the temporal cortex where a tDCS-induced

GABA drop has also been reported (Barron et al., 2016; Koolschijn et al., 2019). Since GABA concentration is higher in grey matter (GM) compared to white matter (WM) (Choi et al., 2006; Jensen et al., 2005), and the GM to WM ratio differed between participants, we also included the GM volume in the MRS voxel in the statistical model.

## METHODS

### Demographics

Data from five studies performed at the Wellcome Centre for Integrative Neuroimaging, University of Oxford were included. Four of these datasets were previously published (Bachtiar et al., 2015, 2018;Barron et al., 2016; Koolschijn et al., 2019). Demographic information for participants in each study are provided in Table 1.

**Table 1:**
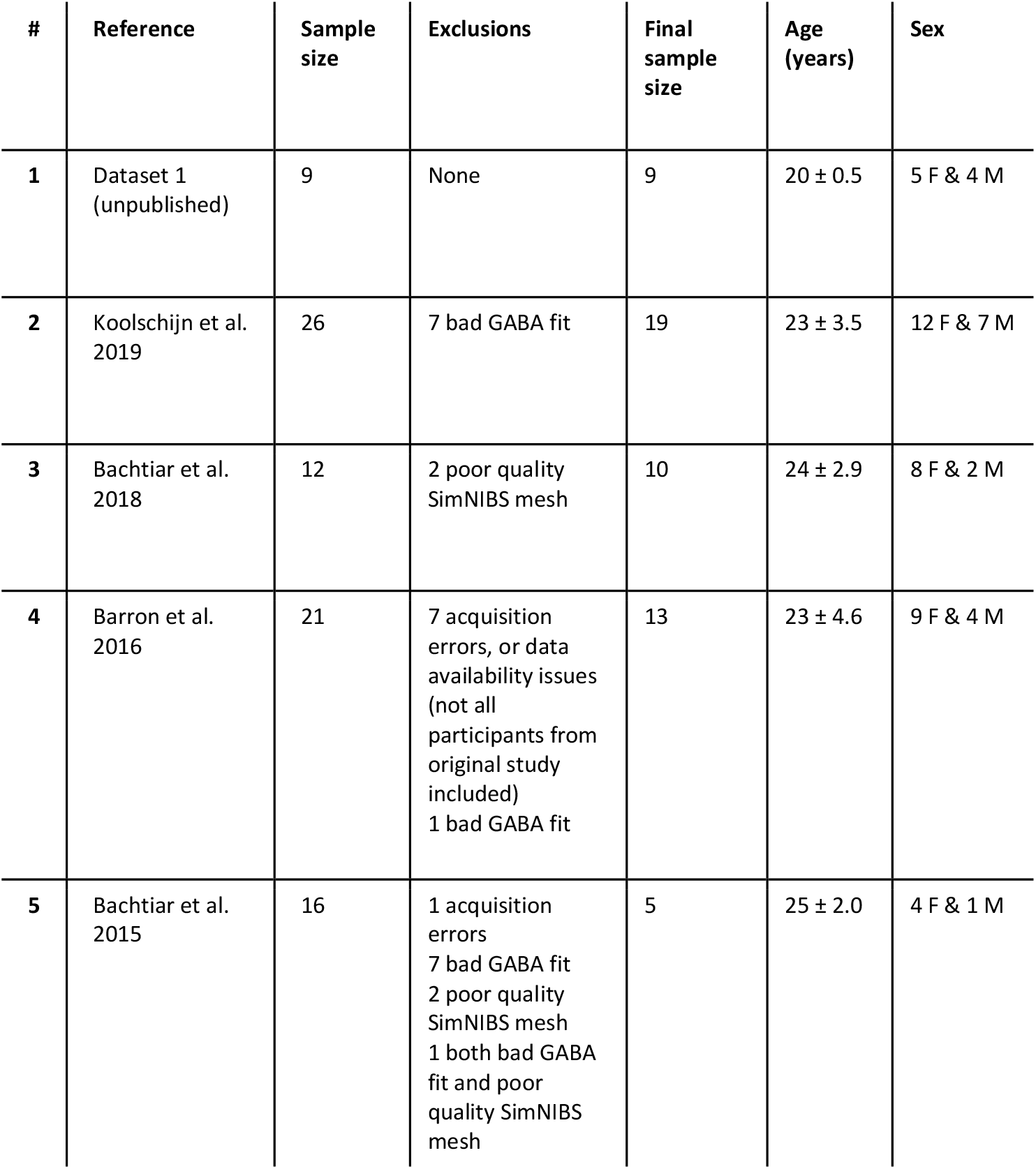
Demographics. Age and sex estimates after accounting for all exclusions.

| # | Reference | Sample size | Exclusions | Final sample size | Age (years) | Sex |
| --- | --- | --- | --- | --- | --- | --- |
| 1 | Dataset 1 (unpublished) | 9 | None | 9 | 20 ± 0.5 | 5 F & 4 M |
| 2 | Koolschijn et al. 2019 | 26 | 7 bad GABA fit | 19 | 23 ± 3.5 | 12 F & 7 M |
| 3 | Bachtiar et al. 2018 | 12 | 2 poor quality SimNIBS mesh | 10 | 24 ± 2.9 | 8 F & 2 M |
| 4 | Barron et al. 2016 | 21 | 7 acquisition errors, or data availability issues (not all participants from original study included)<br>1 bad GABA fit | 13 | 23 ± 4.6 | 9 F & 4 M |
| 5 | Bachtiar et al. 2015 | 16 | 1 acquisition errors<br>7 bad GABA fit<br>2 poor quality SimNIBS mesh<br>1 both bad GABA fit and poor quality SimNIBS mesh | 5 | 25 ± 2.0 | 4 F & 1 M |

### tDCS application

Details of tDCS application are provided in table 2 and figure 1. All tDCS was 1mA, and the cathodal electrode was placed over the contralateral supra-orbital ridge in all cases. We only analysed anodal effects, though some of the original studies included both anodal and cathodal stimulation.

**Table 2:** tDCS application details along with acquisition details of the MRS and structural MRI sequences. LOC: lateral occipital complex; SOR: supra-orbital ridge.

| # | Reference | Intensity (mA) | Duration (min) | Anode | Cathode | Electrode dimensions (cm*cm) | Scanner and field strength | MRS Sequence | MRS TR/TE (ms) | Voxel size (cm<sup>3</sup>) | Number of averages | Acquisition duration (min) | Location (MRS) | Sequence (Structural) | TR (ms)/ TE (ms)/ flip angle | Resolution (mm) / Grappa factor |
| --- | --- | --- | --- | --- | --- | --- | --- | --- | --- | --- | --- | --- | --- | --- | --- | --- |
| 1 | Dataset 1 (unpublished) | 1 | 10 | Left M1 | Right SOR | 5*7 | Siemens Magnetom 7T | Semi-LASER | 5000/36 | 2*2*2 | 64 | 5.3 | Primary motor cortex | MPRAGE | 2200/2.82/7° | 1x1x1/2 |
| 2 | Koolschijn et al. 2019 | 1 | 20 | Right LOC (temporal cortex) | Left SOR | 5*7 | Siemens Magnetom 7T | Semi-LASER | 5000-6000/36 | 2*2*2 | 65 – 130 | 10 | Temporal cortex | MPRAGE | 2200/2.96/- | 0.7x0.7x0.7/- |
| 3 | Bachtiar et al. 2018 | 1 | 10 | Left M1 | Right SOR | 5*7 | MRS: Siemens Magnetom 7T<br>Structural: Siemens Verio 3T | Two-voxel interleaved semi-LASER<sup>1</sup> | 7000/30 | 2*2*2 | 64 per voxel | 12 | Primary motor cortex | Multigradient echo | 2530/1.79,3.65,5.51,7.37/- | 1x1x1/- |
| 4 | Barron et al. 2016 | 1 | 20 | Right LOC (temporal cortex) | Left SOR | 5*7 | Siemens Magnetom 7T | Semi-LASER | 5000-6000/36 | 2*2*2 | 96 - 128 | 10 | Temporal cortex | MPRAGE | 2200/2.96/- | 0.7x0.7x0.7/- |
| 5 | Bachtiar et al. 2015 | 1 | 20 | Left M1 | Right SOR | 5*7 | Siemens Verio 3T | MEGA-PRESS | 2000/68 | 2*2*2 | 144 * 3 | 13.5 | Primary motor cortex | MPRAGE | 2040/5/8° | 1x1x1/- |

**Figure 1:**
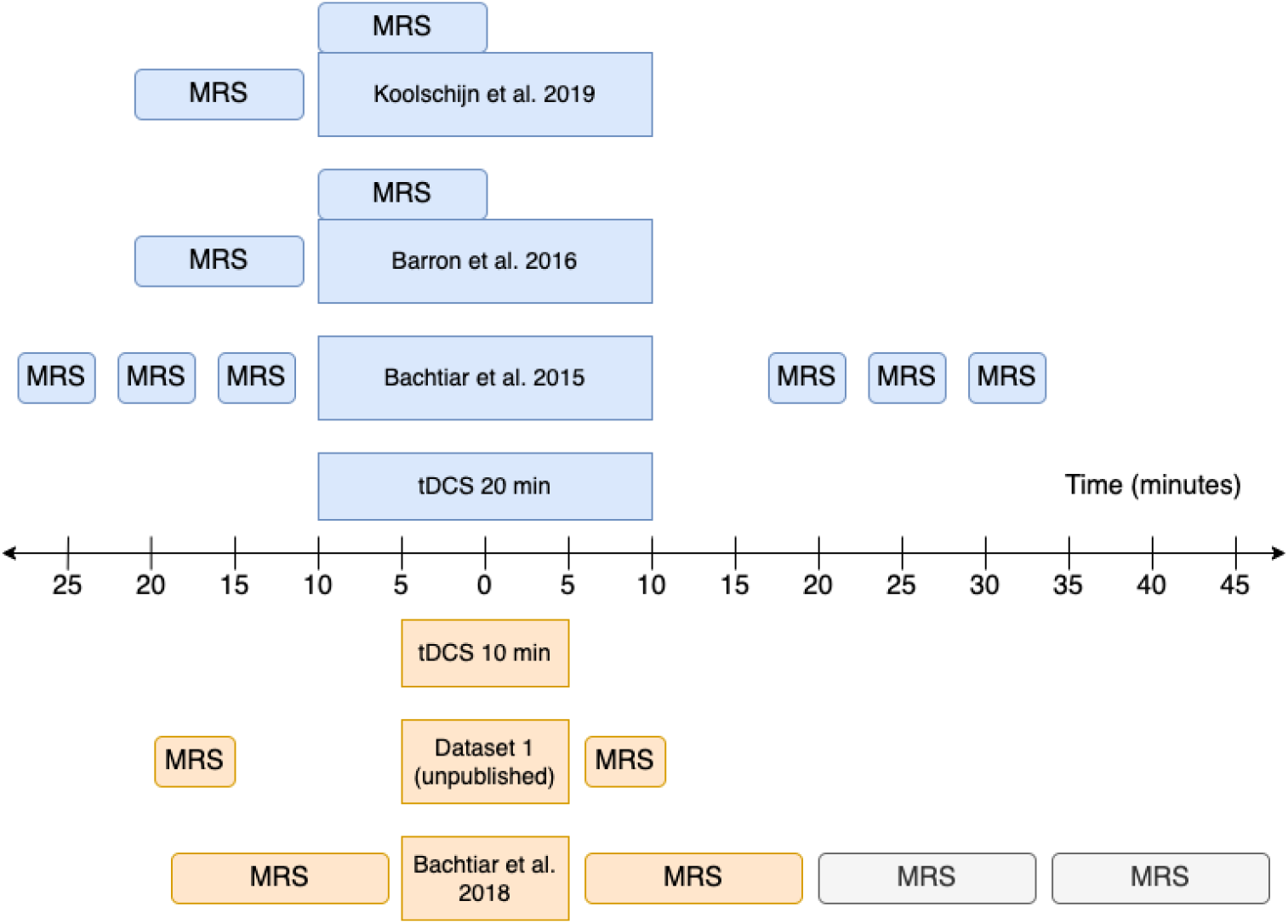
MRS acquisition timelines. Greyed out acquisition blocks were not included in the analysis. All studies used anodal tDCS. Studies using 20 mins tDCS are shown in blue (top) and those using 10 mins tDCS are shown in orange (bottom). For other study parameters see Table 12.

### MRS acquisition

MRS data were acquired at either 3T using a MEGA-PRESS sequence, or at 7T using a semi-LASER sequence. Further details of the acquisition protocols including TR/TE, voxel size and acquisition duration, are provided in table 2 and figure 1.

### MRS analysis

#### Semi-LASER data

Standard preprocessing was applied (Near et al., 2021), including eddy current correction, phasing of spectra and residual water removal using Hankel–Lanczos singular value decomposition (HLSVD). Coil combination used the complex weights of the water unsuppressed reference data calculated using Brown’s method (Brown, 2004). Neurochemicals were quantified using linear combination fitting in LCModel (Provencher, 2001). Fitting used basis spectra containing 19 metabolites (L-Alanine, Ascorbate, Aspartate, Glycerophosphocholine, Phosphocholine, Creatine, Phosphocreatine, γ-Aminobutyric Acid, Glucose, Glutamine, Glutamate, Glutathione, myo-Inositol, L-Lactate, N-Acetylaspartate, N-Acetylaspartylglutamate, Phosphorylethanolamine, scyllo-Inositol, Taurine) and an empirically measured macromolecular basis spectrum. No concentration ratio priors were applied (LCModel parameter *NRATIO* was set to “7”). Metabolite fits with absolute pairwise correlation coefficients above 0.5 were combined. Metabolite concentrations were expressed as a ratio to total-creatine (tCr; Creatine+Phosphocreatine), not corrected for GM concentration. The following criteria were applied for excluding poor quality spectra: Cramér-Rao lower bounds (CRLB) > 50% and/or LCModel-reported-SNR < 30. Additionally, GABA:tCr ratios < 0 or > 1, were excluded. MRS data quality information for included participants is provided in supplementary table 1.

#### MEGA-PRESS data

Pre-processing and fitting was achieved using a combined FID-A (Simpson et al., 2017, p.) and Gannet (Edden et al., 2014) pipeline. Specifically, the data were concatenated and pre-processed using FID-A (*run_megaoressproc_auto* script) before being zero-padded and filtered (3 Hz line broadening) to match Gannet’s pre-processing. Finally, it was fitted and quantified using Gannet (*GannetFit*, *GannetCoRegister*, *GannetSegment*, *GannetQuantify*). The following criteria were applied to excluding poor quality spectra: NAA linewidth > 10 Hz, and a GABA:Cr ratio of < 0 or > 1. MRS data quality information for included participants is provided in supplementary table 1.

### E-field modelling

The head models were built using an in-house implementation combining a new segmentation approach (charm) (Puonti, Van Leemput, et al., 2020) with the standard headreco pipeline (Saturnino, Puonti, et al.,2019) in SimNIBS version 3.2. Specifically, each subject’s MRI scan (see table 2 for acquisition details) was processed with both headreco and charm, and a fused head segmentation was generated by combining the brain tissue segmentations, and grey matter surfaces, from headreco with the extra-cerebral segmentations from charm. Finally, a finite element (FEM) mesh was generated, including representations of the scalp, skull, cerebrospinal fluid (CSF), GM, and WM, which was subsequently used for the electric field simulations. All segmentations were manually inspected, and edited when needed, to ensure that the head models were accurate. The mean and 95^th^ percentile of the E-field magnitude as well as the components of the E-field normal and tangential to the grey matter surface were estimated. Separate statistical analyses were run for the E-field within the MRS voxel, and the E-field over the whole cortical surface. For the voxel analysis, the E-field values within the MRS voxel were extracted in individual space, while for the whole brain analyses the E-field values were analysed in fsaverage space. See figures 2 & 3 for the E-field components within the MRS voxel, and figures 4 & 5 for the E-field components over the whole cortex. In figures 2 & 3 the soft voxel mask, defined as the overlap fraction of the MRS voxel mask across subjects, was thresholded at 10% to include most of the area covered by the MRS voxel in individual subjects.

**Figure 2:**
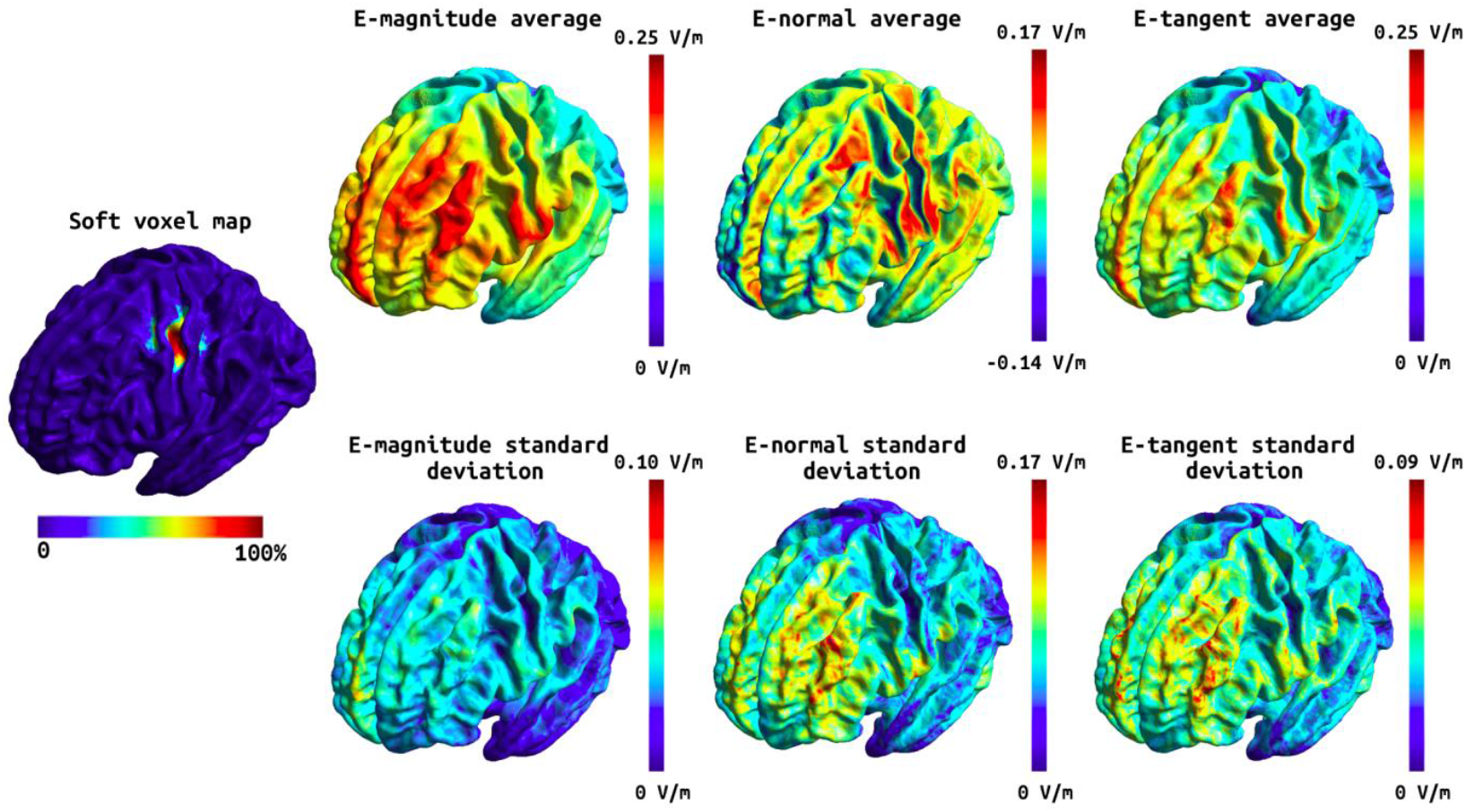
Soft voxel map (leftmost) and the mean and standard deviation of the E-field components over the subjects in fsaverage space for the tDCS stimulation targeting the M1. First column: the mean E-field magnitude (top) and its standard deviation (bottom). Second column: the mean E-field normal component (top) and its standard deviation (bottom). Third column: the mean E-field tangential component (top) and its standard deviation (bottom). ote that the normal component has directionality where positive (red) values denote currents flowing into the cortex and negative (blue) values denote currents flowing out of the cortex.

**Figure 3:**
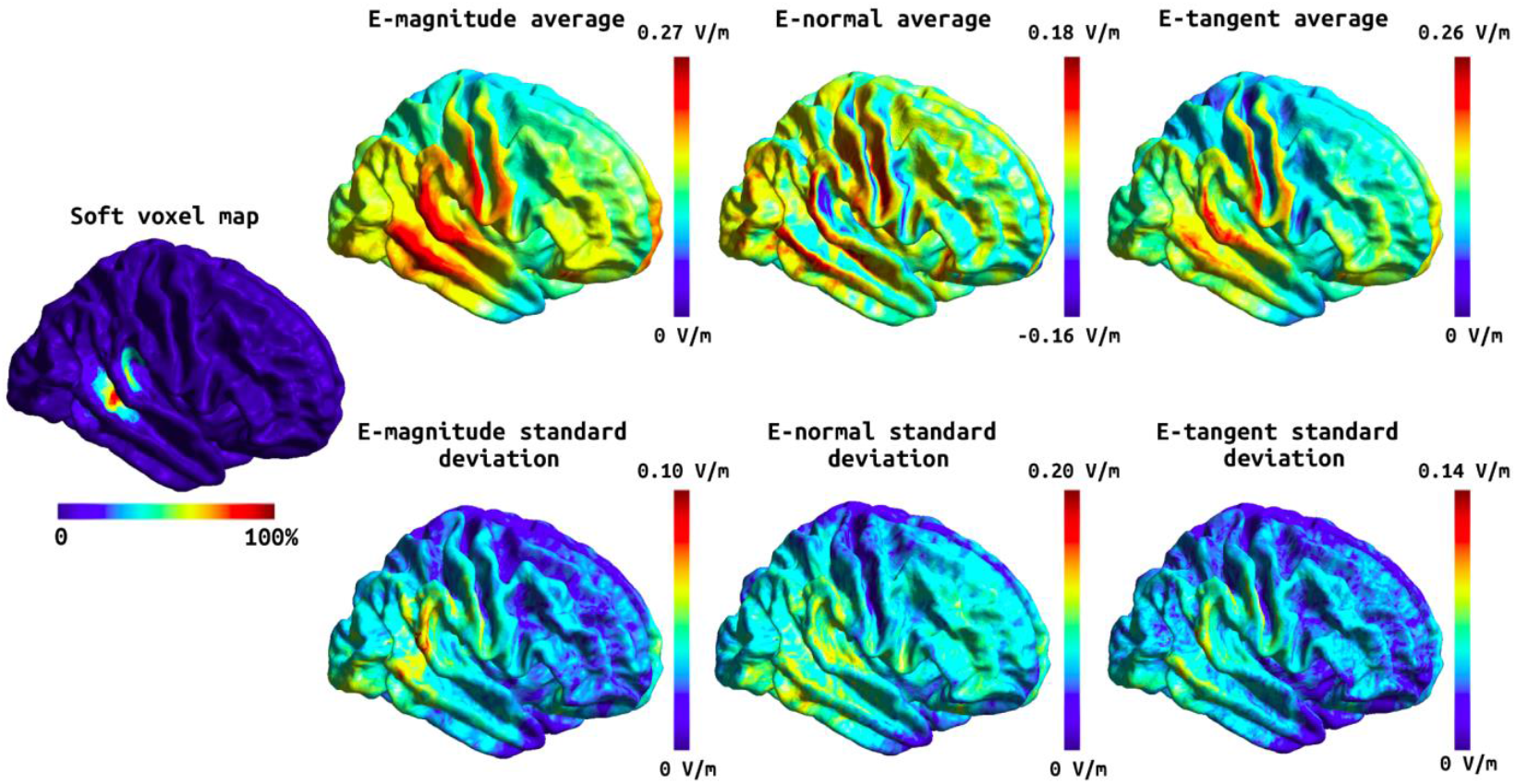
Soft voxel map (leftmost) and the mean and standard deviation of the E-field components over the whole cortex in fsaverage space for the tDCS stimulation targeting the temporal cortex. First column: the mean E-field magnitude (top) and its standard deviation (bottom). Second column: the mean E-field normal component (top) and its standard deviation (bottom). Third column: the mean E-field tangential component (top) and its standard deviation (bottom). Note that the normal component has directionality where positive (red) values denote currents flowing into the cortex and negative (blue) values denote currents flowing out of the cortex.

**Figure 4:**
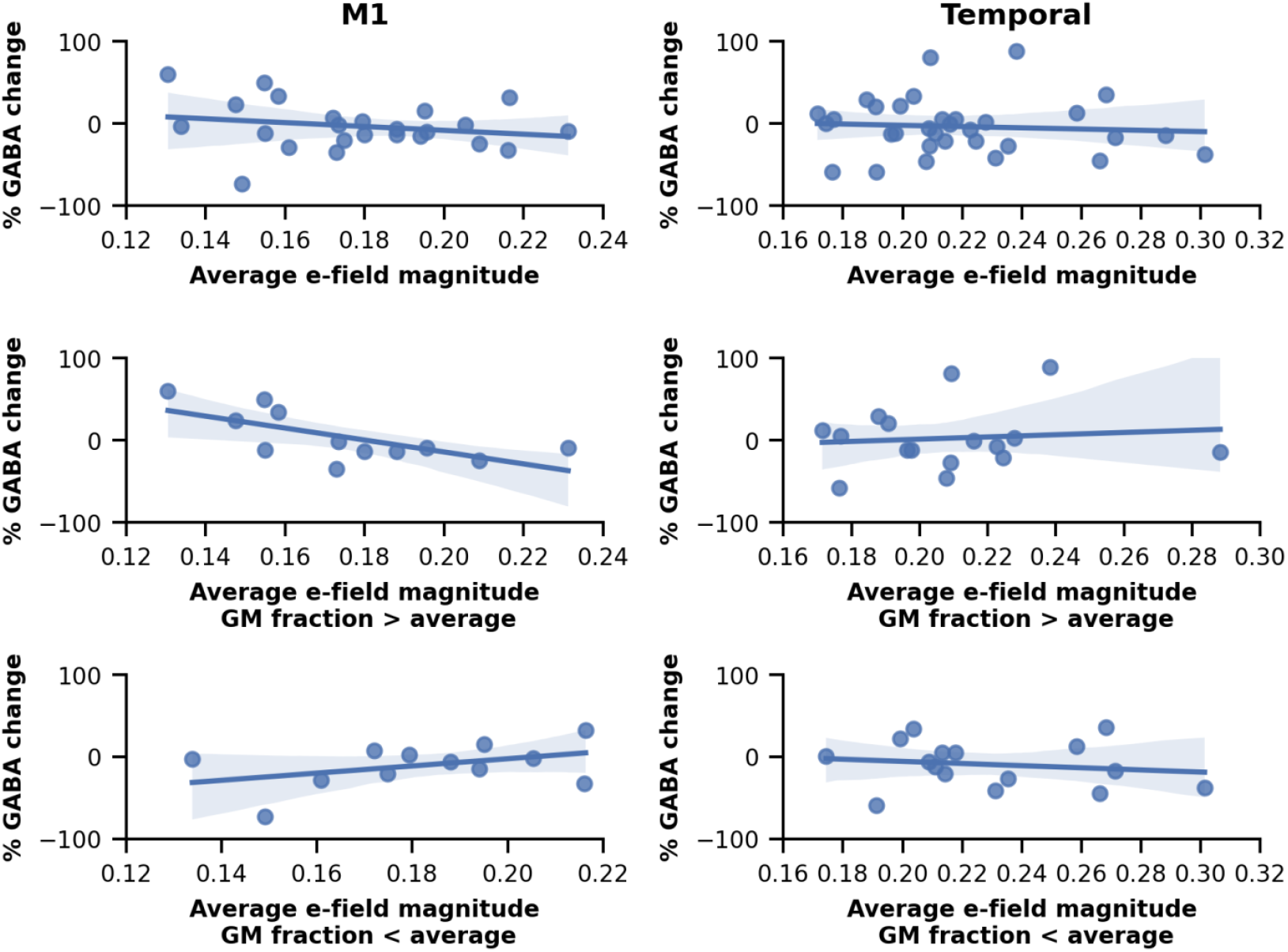
Percent GABA change plotted against the average electric field magnitude in the M1 MRS voxel (left) and in the temporal MRS voxel (right). The two lower rows show the relation between percent GABA change and the electric field magnitude when the subjects are split according to the grey matter fraction in the MRS voxel (above or below the average).

### Statistical analysis

#### MRS voxel

Statistical analyses were conducted using R (RCoreTeam2013). Percentage change in GABA was calculated, and we used the robust outlier detection method based on the adjusted box-plot rule within the MATLAB toolbox to identify outliers (Pernet et al., 2013). One outlier was identified which was then excluded from further analysis. To test whether induced E-field was correlated with tDCS-induced GABA change, and whether this effect depended on GM volume in the area of interest, we used linear mixed-effects (LME) models. Due to well documented problems with accurately modelling physiological processes on percentage change values (Curran-Everett & Williams, 2016), we subjected the raw GABA values to change analysis using LMEs (Curran-Everett & Williams, 2015). To that end, we constructed an LME model of GABA using the R package lme4 (Bates et al. 2010) and included timepoint (pre, post), E-field, and grey matter volume as fixed effects, as well as a three-way interaction effect of time * E-field * grey matter volume as the effect of interest (Equation 1). All two-way interactions are included in the model by default (time * E-field, time * grey matter volume, and E-field * grey matter volume). We allowed intercepts for different subjects to vary to account for covarying residuals within subjects. Initially, we allowed intercepts to also vary for different studies, to account for co-varying residuals within studies. However, because the variance captured by the random effect of study was approximately zero, and likelihood ratio testing indicated that the random effect of study was not significant, it was dropped from the model. Finally, p-values were obtained using the anova function from the lmerTest package, which uses the Satterthwaite’s method for denominator degrees-of-freedom and F-statistic (Kuznetsova et al.,2017). Mean ± SD are presented throughout. A total of twelve LME models were run, one for each E-field variable (mean and 95th percentile of magnitude, normal and tangential components), and separately for the M1 and temporal data.

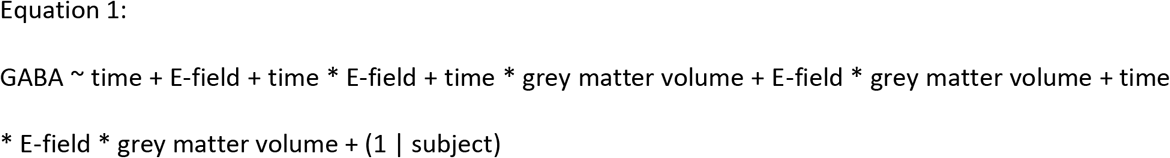

In case of a significant three-way interaction, posthoc tests were run using the TestIntercations functions from the phia package. Specifically, we tested the interactions between E-field and time, at the 25th and 75th percentile grey matter volume values. We performed a full statistical model that included all main effects and interactions by default. For transparency, we include all these results in the supplementary materials.

#### Whole brain analysis

In addition to the MRS voxel analysis, we performed a whole brain cortical surface analysis to determine whether tDCS-induced change in GABA was correlated with different E-field components outside the MRS voxel. To this end, we used the Permutation Analysis of Linear Models (PALM, version alpha119) software tool (Winkler et al., 2014) to perform the analysis in Matlab (version 9.3.0.713579 (R2017b), The Mathworks Inc, Natick, Massachusetts). Specifically, we ran a separate linear model for each of the three E-field components (magnitude, normal, tangent), and stimulation site (M1, temporal), using the GABA change as an independent variable. The effect of study site (three sites for M1 and two for temporal, see table 3) was included as a nuisance variable, and permutations were performed only within the data from each site. Two contrasts were defined: one corresponding to a positive linear relationship between the tDCS-induced GABA change and the E-field component and the other corresponding to a negative linear relationship. One thousand permutations were performed for each model (six models in total), with the shuffling restricted to the study specific variance groups (option -vg auto) (Winkler et al., 2015), using threshold-free cluster enhancement (option -T) (Smith & Nichols, 2009) and tail approximation (option - accel tail) (Winkler et al., 2016) to reduce execution time. The final whole-cortex maps report the family-wise error corrected p-values for the Aspin–Welch’s *v*-statistic.

## RESULTS

A number of variables arise from the E-field modelling. Of these, we have chosen to focus on the magnitude of the E-field, which has previously been shown to be associated with GABA change (Antonenko et al., 2019), as our variable of interest. However, this choice is not consistent across the literature, hence we have also included other commonly used metrics to aid comparison with the existing literature. All E-field variables for the M1 and temporal data are shown in Figures 2 and 3.

### tDCS induced a decrease in GABA in M1

We first wanted to determine whether we could replicate the previously-reported tDCS-induced decrease in GABA in M1. We demonstrated a significantly lower GABA after tDCS compared with before (Pre: 0.29 ± 0.12, Post: 0.27 ± 0.14, Main Effect of time t(24) = −3.35, p = 0.003; suppl. figure 1).

### Change in GABA is related to E-field in M1

Having established across the group that anodal tDCS led to a decrease in GABA in M1, we then went on to investigate whether the tDCS-induced decrease in GABA was related to the calculated E-field in the M1 voxel on a subject-by-subject basis. There was a significant interaction between time and mean E-field magnitude (t(24) = 3.24, p = 0.003) indicating, as hypothesised, that higher E-fields were associated with greater tDCS-induced decreases in GABA (Fig. 4). This effect was consistent across all E-field variables except the average of the normal component (Suppl. fig. 2, suppl. table 2).

### Proportion of GM in the MRS voxel moderates the relationship between E-field and change in GABA in M1

In M1, a significant three-way interaction (t(24) = −3.55, p = 0.002) between time, mean E-field magnitude and the proportion of GM in the voxel, revealed that the association between E-field and GABA change was moderated by the GM content in the MRS voxel. Post-hoc tests demonstrated that the association between E-field and GABA change was present only in voxels with a relatively high GM content (at 75% percentile, Chisq(1) = 12.91, p < 0.001), but not in those with a relatively low GM content (at 25% percentile Chisq(1) = 1.78, p=0.128) (Fig. 4). Again, this effect was consistent across all E-field variables except the average of the normal component (Suppl. Figs. 3 and 4, suppl. table 3). Afterwards, we also ran the reduced linear models, adding the predictors and their interactions one-by-one, to check if the three-way interaction is necessary for obtaining a statistically significant relation between GABA and the E-field. It was only after the GM proportion was added that statistical significance was reached.

### Similar relationships are not observed in temporal cortex

Finally, to determine whether E-field and grey matter volume explain tDCS-induced GABA decreases outside M1, we analysed data from a voxel in the temporal cortex. tDCS has previously been reported to decrease GABA in this region (Barron et al., 2016; Koolschijn et al., 2019). However, unlike M1, we did not observe a significant relationship between E-field and tDCS-induced GABA drop (Fig. 4 and suppl. fig. 5), even when accounting for volume of GM in the voxel (Suppl. figs. 6 and 7, suppl. table 4)

### Whole-brain statistical analysis

Finally, we explored whether there were any regions in the brain where E-field significantly related to the tDCS-induced GABA decrease within our MRS voxel. We found no significant clusters for either the M1 or the temporal stimulation setup.

## DISCUSSION

This study was performed to address the hypothesis that a significant amount of the inter-subject variability in the anodal tDCS-induced decreases in GABA could be explained by the individual E-field within the stimulated region. In line with this hypothesis, we found that tDCS-induced decrease in GABA was associated with the induced E-field in the MRS voxel, supporting the future use of individualised dosing to minimise inter-individual variability in tDCS effects. Further, we showed that this association is more complex than previously demonstrated: the relationship between GABA decrease, and E-field was only demonstrated in participants who had a relatively high volume of grey matter in their MRS voxel. In addition, while a significant correlation between GABA decrease, and E-field was demonstrated in M1, this was not present in the temporal cortex.

### Anodal tDCS-induced M1 GABA change is associated with the induced E-field

One of the factors limiting the therapeutic potential of tDCS is the high inter-subject variability of behavioural effects. It has been suggested that this might, at least in part, be due to the use of a standard extracranial current intensity across participants, which will inevitably lead to variability in the E-field applied in the cortex (Abellaneda-Pérez et al., 2020; Kim et al., 2014; Laakso et al., 2015; Mezger et al.,2021; Soleimani et al., 2021). Supporting this theory, tDCS-induced behavioural outcomes such as improvements in working memory have been related to the applied E-field (Albizu et al., 2020). Some studies have gone further and tried to link neurophysiological changes to applied E-field, showing that the magnitude of tDCS-induced corticospinal excitability changes (Laakso et al. 2019), and decreases in both glutamate (Mezger et al., 2021) and GABA (Antonenko et al., 2019) are related to the intensity of the simulated current in the stimulated region. However, the anatomical location of the relationship between neurophysiological changes and E-field has not been consistently demonstrated, even in studies investigating similar brain regions and stimulation montages. For example, Antonenko and colleagues showed no significant correlation between E-field in their M1 MRS voxel and tDCS-induced GABA decrease, although they demonstrated a significant relationship between M1 GABA change and the E-field in a pre-central gyrus cluster that was functionally connected to the MRS voxel during stimulation (Antonenko et al., 2019). It is not clear why our results are not perfectly in line with those of Antonenko and colleagues, but it may be due to methodological differences between the two studies. Here, we used a large ROI, which reflected the entire MRS voxel, unlike the smaller spherical ROI employed by Antonenko and colleagues to represent the E-field in the MRS voxel.

Here, we investigated the relationship between tDCS-induced GABA and E-field both with an ROI approach, using the MRS voxel from which we quantified GABA, and across the whole brain. In our ROI-based approach we demonstrated that the tDCS-induced decrease in GABA was correlated with mean E-field within the M1 MRS voxel; a relationship mediated by the volume of grey matter in the voxel. Using single voxel approaches, we are only able to quantify tDCS-induced GABA change in a single MRS voxel, and therefore inherently bias our results to this location. Though our whole brain analysis did not show any significant effects, it is possible that our exploratory analysis was not adequately powered to address this question. tDCS is known to have remote effects in areas that are anatomically and/or functionally connected to the region where the E-field is quantified (Mezger et al., 2021), and E-fields outside the MRS voxel may also influence physiological effects captured within the voxel. Further studies are required to examine whether the effective use of tDCS may require optimization of off-target effects based on individual anatomy, in addition to individualised dosing.

### Anodal tDCS-induced GABA change is more associated with E-field in the MRS voxel in subjects with high GM volumes

Both animal (Fahn & Ccote, 1968) and human studies (Choi et al., 2006; Jensen et al., 2005) suggest that GABA concentration is higher in GM compared to WM, and in line with this, in our data GABA estimates were higher in M1 MRS voxels with higher grey matter volume (Suppl. fig. 8). Therefore, in participants who had very small amounts of GM within their MRS voxel, any effects on GABA concentration may be too small to detect, or may suffer from a floor effect. This does not rule out the possibility that tDCS actually induced a GABA change, but instead reflects a methodological limitation in our ability to detect any such changes. It is possible that an association between E-field and GABA change would become evident in these participants if the MRS voxel was shifted to include more GM, or through use of MRSI techniques.

### Magnitude, normal and tangential components of E-field

In addition to the magnitude, the orientation of the E-field relative to the cortical surface and consequently the neurons, can also influence the effectiveness of tDCS (Radman et al., 2009; Rawji et al.,2018). *In vitro* data suggests that E-fields parallel to the main axis of a neuron are more effective than perpendicular E-fields for polarising neuronal membranes (Radman et al., 2009). Since pyramidal neurons are oriented perpendicular to the cortical surface, it would follow that that E-fields perpendicular (normal) to the cortical surface would have a greater neurophysiological effect. Additionally, in theory, the symmetric dendritic morphology of interneurons makes them more difficult to polarise (Radman et al.,2009). Rahman et al. (Rahman et al., 2013), however, showed that the M1-supraorbital montage generates higher tangential, compared to normal E-fields, and their data suggest that tangential fields can acutely modify synaptic efficacy through polarisation of axon terminals. Therefore, in theory, both normal and tangential E-field components may contribute to the net tDCS effect, with different neurons and/or cellular sub-components being targeted by each.

However, the relationship between current direction and neurophysiological changes *in vivo* is less clear. Previous human studies have shown that the direction of current flow relative to the central sulcus or a global coordinate system, influences the neurophysiological (Rawji et al., 2018) and behavioural (Albizu et al., 2020) effects of tDCS. However, these studies did not examine the E-field relative to the cortical surface. Laakso et al. (Laakso et al., 2019), did show that the E-field normal component influenced anodal tDCS-induced corticospinal excitability changes, such that high E-fields were associated with a decrease in excitability, and vice-versa. However, they did not test for a relationship between excitability changes and the magnitude of the E-field. Since the magnitude and normal component are highly correlated, this makes interpreting a relationship between normal E-field and excitability changes difficult. Our data does not suggest that either the normal or tangential E-field components are more strongly associated with tDCS-induced change in GABA than the magnitude. Given the strong *in vitro* relationships between current direction and neurophysiological effects, it is not clear why a similarly strong relationship is not seen *in vivo.* Since there are several uncertainties associated with E-field modelling, it is possible that the angle between the cortical surface and the E-field vector is not estimated fully accurately. Physiologically, GABA drop can be mediated by polarisation of GABA interneurons, but polarisation of pyramidal neurons may also influence GABA by altering their interactions with GABA interneurons. Consequently, in theory, both the normal and tangential components could contribute to the overall effects, and using more fine-grained outcomes that can distinguish between the effects on different types of neurons may shed more light on any differential effects of the components.

### Lack of relationship between E-field and GABA changes in the temporal lobe

We focussed on the relationship between E-field and tDCS induced GABA changes in M1 as this is the region which has been most robustly studied. However, to determine whether the relationship identified in M1 reflected a more general property of the cortex, we additionally considered data from studies that had used a voxel placed in the temporal lobe. Here, unlike M1, we did not demonstrate a significant relationship between tDCS-induced GABA changes and E-field within the voxel. There are several potential reasons for this. Firstly, constructing anatomical head models from structural scans acquired at 7T is more challenging than from those at 3T, due to the larger intensity inhomogeneity artefact, so-called bias field, at higher magnetic fields. While the majority of our subjects in the M1 tDCS had 3T structural scans, all the structural scans in the temporal tDCS studies were acquired on a 7T scanner. Additionally, the signal often drops close to the temporal lobes in 7T structural scans due to limited coil coverage. Although dielectric pads were used during scanning and the scans were processed with an aggressive bias field correction approach to minimise these effects, the resulting head models may still be suboptimal in accuracy around the temporal region.

Secondly, all participants in the temporal studies received 20 min of anodal tDCS, compared to only 5/24 participants in the M1 studies. Monte-Silva et al. 2017 found that increasing the length of anodal tDCS from 13 to 26 min led to a change from increased to *decreased* cortical excitability. Other studies have also shown this reversal of the classic anodal tDCS effect at longer durations and higher intensities (Batsikadze et al., 2013; Hassanzahraee et al., 2020; Simis et al., 2013). This nonlinearity has been suggested to reflect homeostatic metaplasticity mechanisms, which aim to maintain excitation/ inhibition balance, and may be initiated when a certain combination of intensity and duration is exceeded. It is therefore possible that in the temporal studies, a subset of participants displayed homeostatic metaplasticity, driven by a combination of the relatively longer duration and variable E-fields i.e., effective intensity, leading to a reversal of tDCS effects. Consequently, any GABA changes may not be simply related to the applied E-field alone.

### Individualised dosing

Our data, together with other studies (Albizu et al., 2020; Antonenko et al., 2019; Laakso et al., 2019;Mezger et al., 2021), strongly support the use of individualised dosing based on a priori E-field modelling, and algorithms to estimate the extracranial intensity required to achieve a target intracranial E-field have already been established (Caulfield et al., 2020; Evans et al., 2020; Saturnino, Siebner, et al., 2019). In the absence of MRIs, the transcranial electrical stimulation (TES) motor threshold (Caulfield et al., 2020) or head circumference (Antonenko et al., 2021) have been suggested as proxies for the E-field, making individualised dosing more clinically feasible.

One challenge that remains is to determine the optimal or necessary E-field required to achieve a given neurophysiological or behavioural change. In our data, the highest E-field observed in M1 was just over 0.31 V/m (95% percentile E-field magnitude), and a GABA drop was seen in many participants with even lower E-fields. In a subset of our M1 data (n=9), the GABA drop was even accompanied by improvement in a temporal order judgement task. Much larger sample sizes are necessary to establish the minimal E-field necessary to achieve a behaviourally relevant GABA drop, that is over and above any natural physiological fluctuations and measurement error. This threshold will likely vary depending on the neurophysiological and behavioural outcome of interest, and any non-linear effects of intensity must also be considered (Mosayebi Samani et al., 2019). Data from *in vitro* and animal studies (Filmer et al., 2020) will be helpful for elucidating the underlying cellular and network level effects at these effective E-fields.

### Conclusions

We show that in M1, E-field in the MRS voxel is related to the GABA drop, adding to the accumulating evidence that supports individualised dosing of tDCS. The interaction with GM volume within the MRS voxel emphasises the need to appropriately choose and evaluate any outcome measures which we expect to be related to E-field. While we did not find a similar association in the temporal region, given the challenges of modelling the E-field in this region and possible homeostatic metaplastic effects, such an association cannot be ruled out.

## Acknowledgements

The research was supported by the National Institute for Health Research (NIHR) Oxford Biomedical Research Centre and the NIHR Oxford Health Biomedical Research Centre. The Wellcome Centre for Integrative Neuroimaging is supported by core funding from the Wellcome Trust (203139/Z/16/Z). TN was supported by a NIHR Oxford Biomedical Research Centre Small Grant Award. HB is supported by the Medical Research Council (MRC) UK (MC_UU_00003/4). CJS holds a Sir Henry Dale Fellowship, funded by the Wellcome Trust and the Royal Society (102584/Z/13/Z). This project has received funding from the European Union’s Horizon 2020 research and innovation programme under grant agreement No. 731827 (STIPED). The results and conclusions in this article present the authors’ own views and do not reflect those of the EU Commission. The study was also supported by the Lundbeck foundation (grant R313-2019-622), and by the NIH (Grant No. 1RF1MH117428-01A1).

## Data statement

MRS and structural MRI data can be made available on request.

