## Supplementary Material for "tDCS induced GABA change is associated with the simulated electric field in M1, an effect mediated by grey matter volume in the MRS voxel"

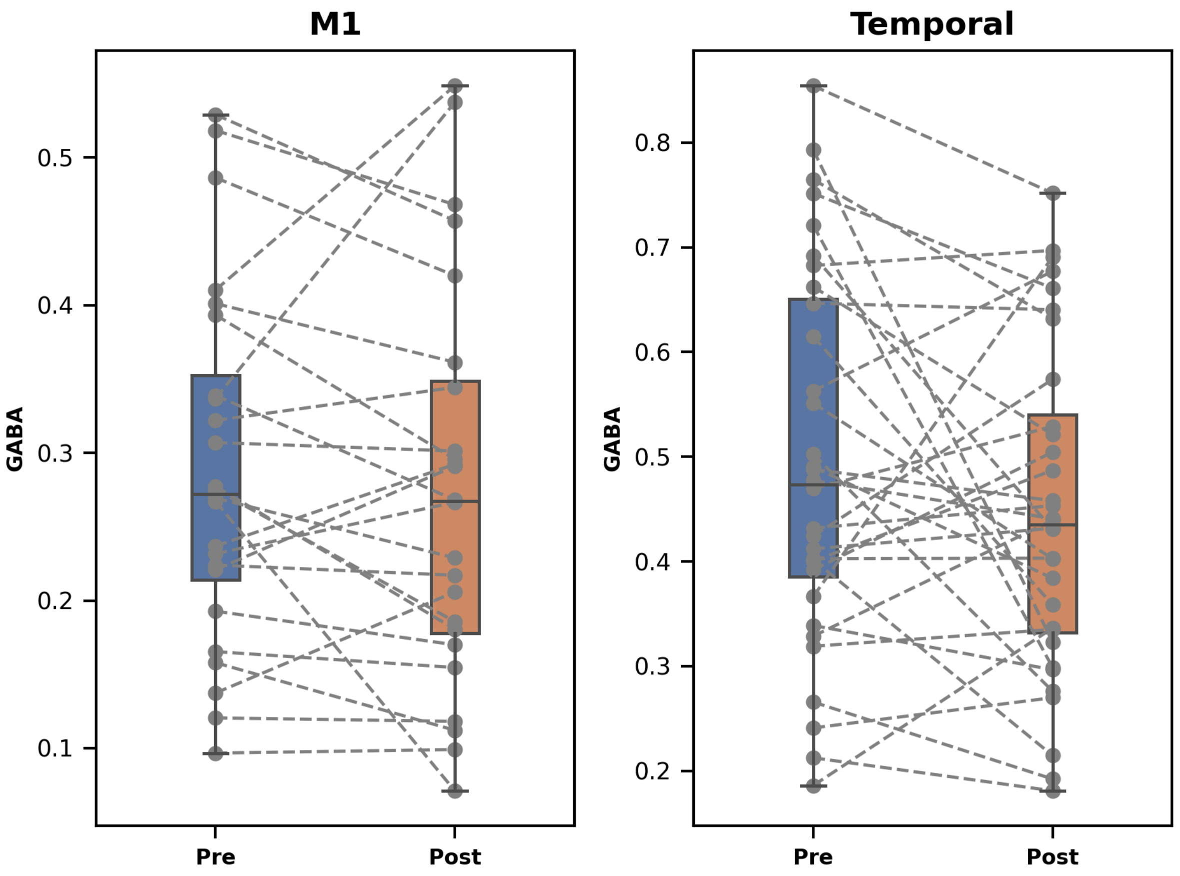


*Supplementary figure 1: Pre and post GABA values for the motor region (left) and the temporal region (right).*

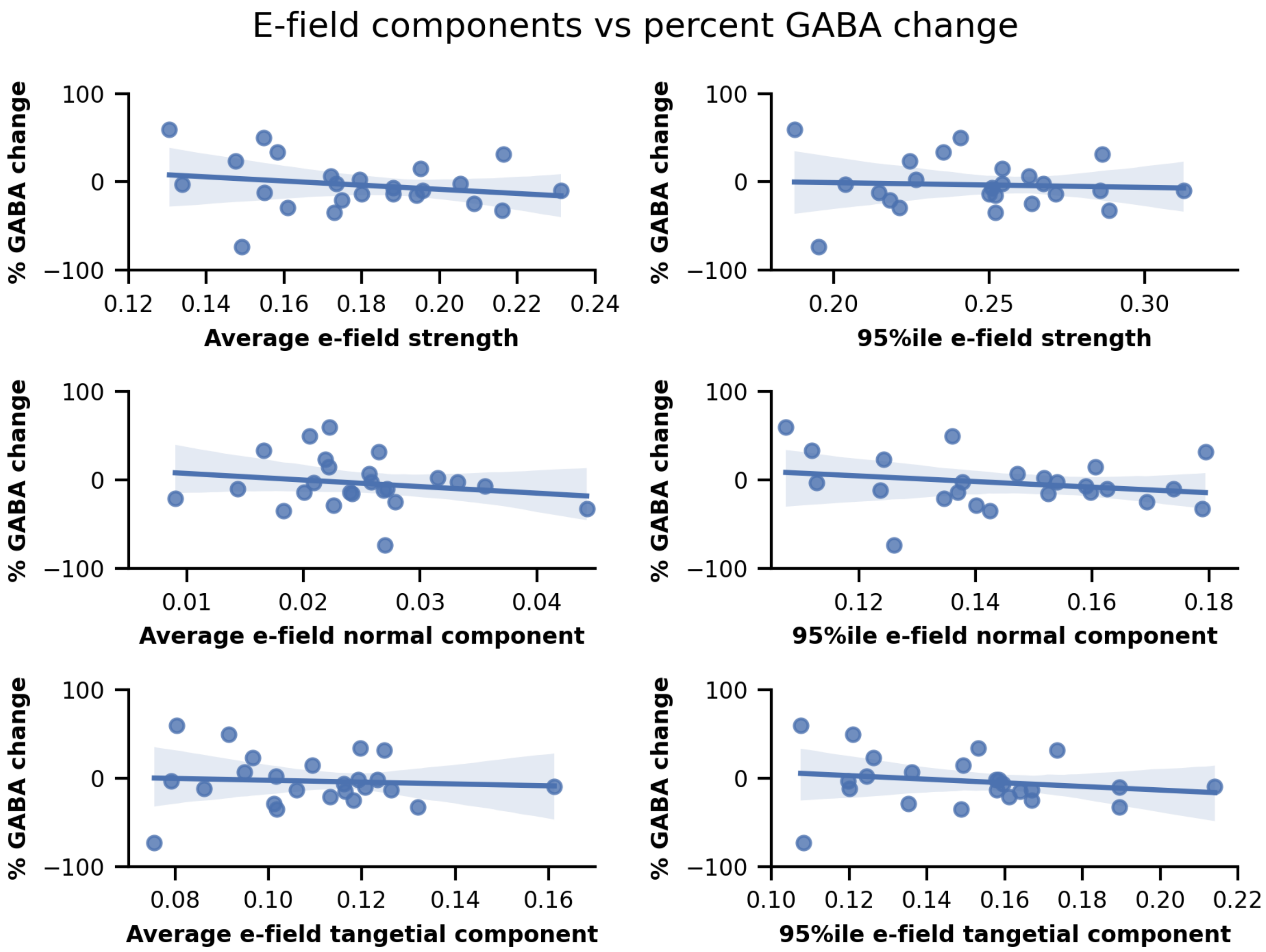


*Supplementary figure 2: Association between all the E-field estimates (magnitude, normal and tangent; mean and 95th percentile), in the M1 MRS voxel, and the percent GABA change.*

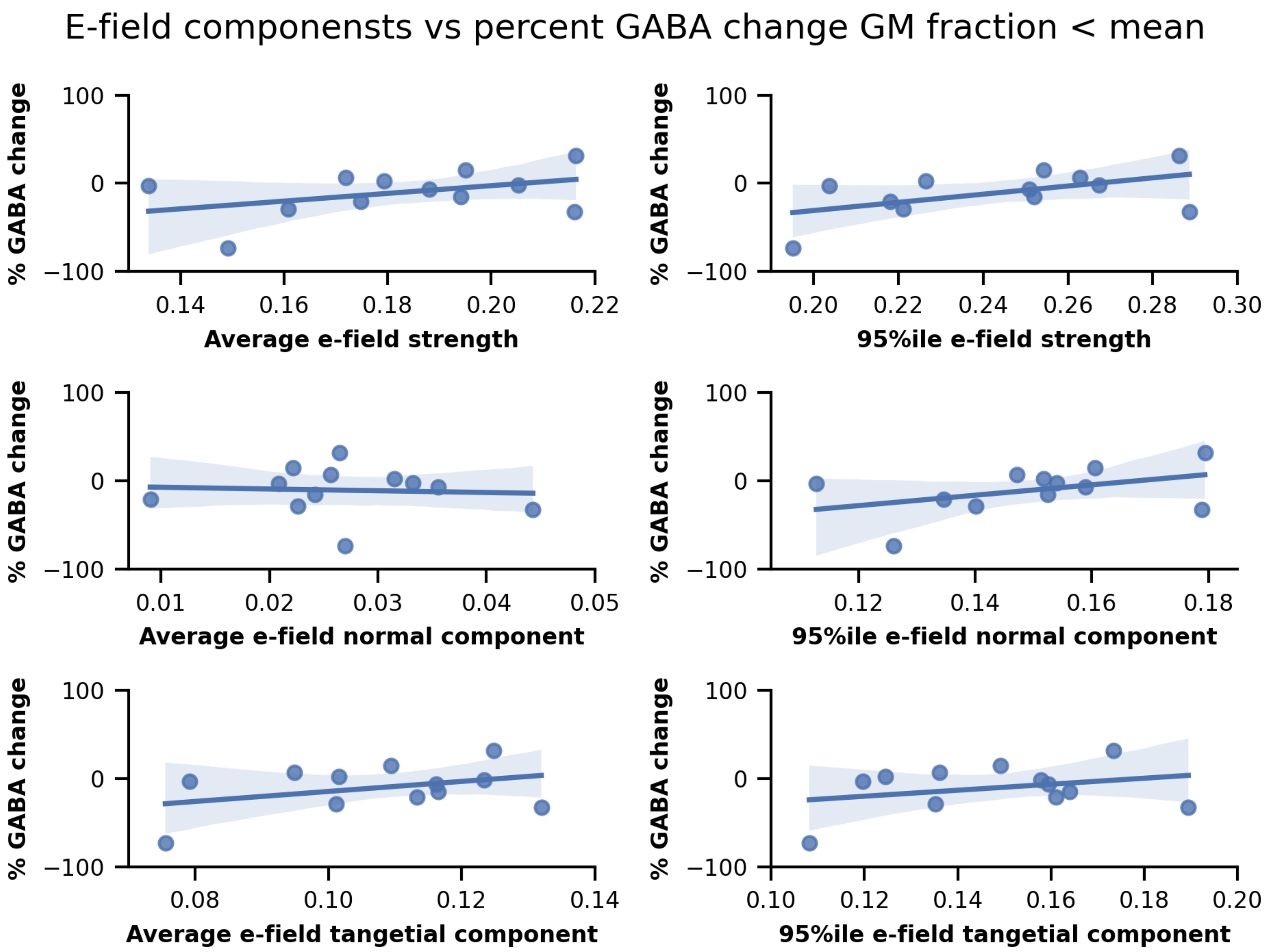


*Supplementary figure 3: Association between all the E-field estimates (magnitude, normal and tangent; mean and 95th percentile), in the M1 MRS voxel, and the percent GABA change, only for participants with grey matter partial volume estimates lower than the mean.*

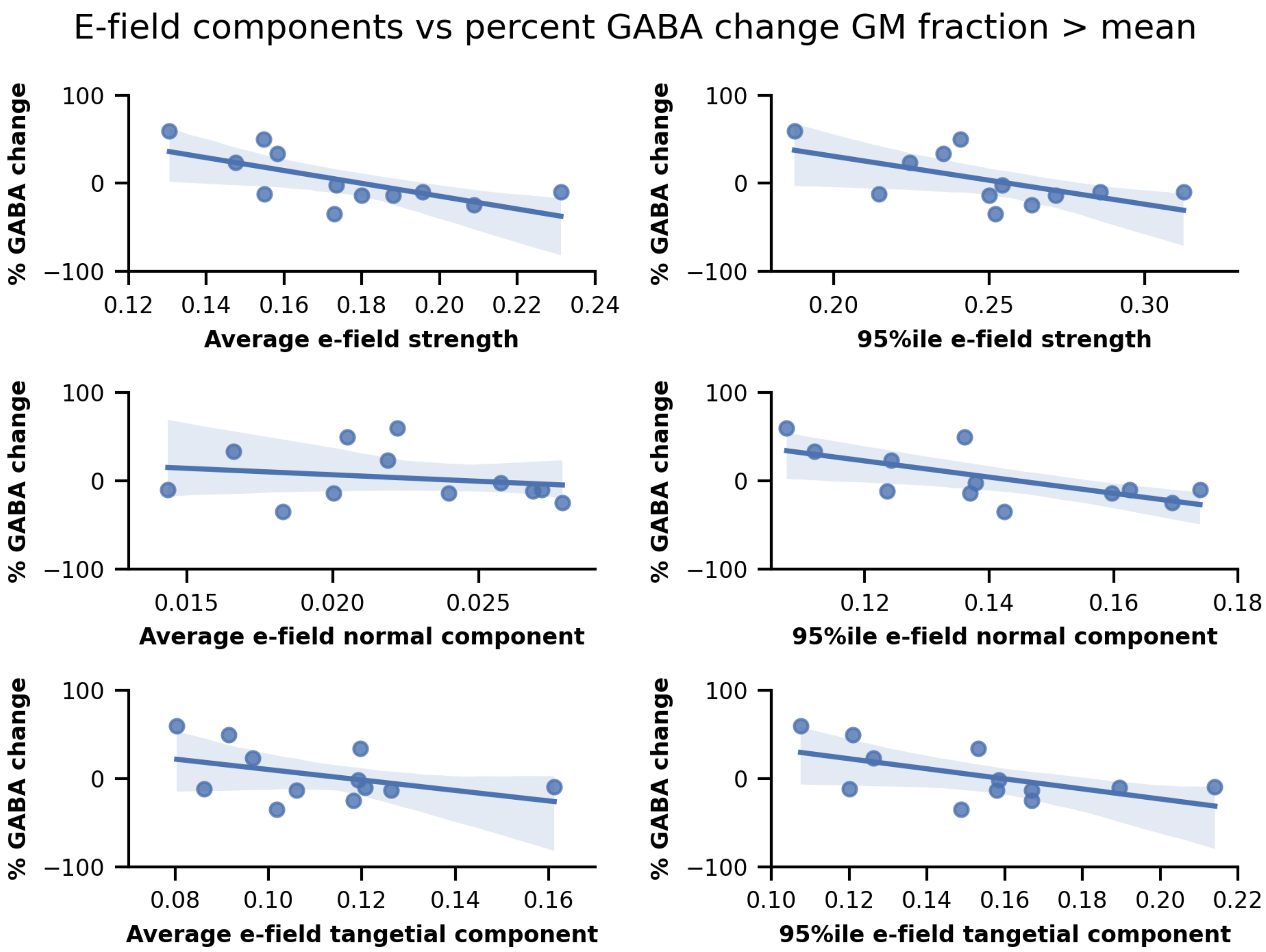


*Supplementary figure 4: Association between all the E-field estimates (magnitude, normal and tangent; mean and 95th percentile), in the M1 MRS voxel, and the percent GABA change, only for participants with grey matter partial volume estimates greater than the mean.*

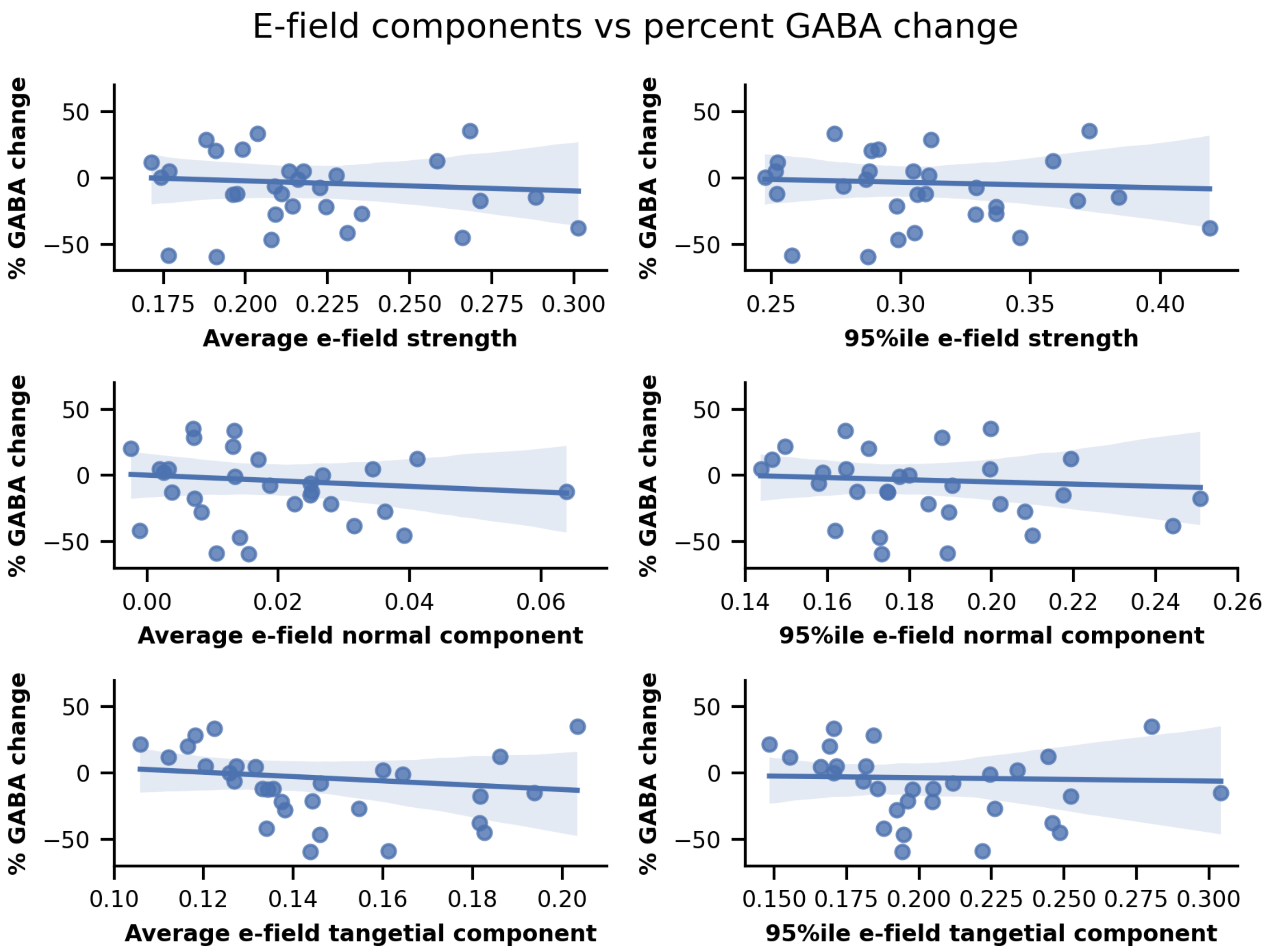


*Supplementary figure 5: Association between all the E-field estimates (magnitude, normal and tangent; mean and 95th percentile), in the temporal MRS voxel, and the percent GABA change.*

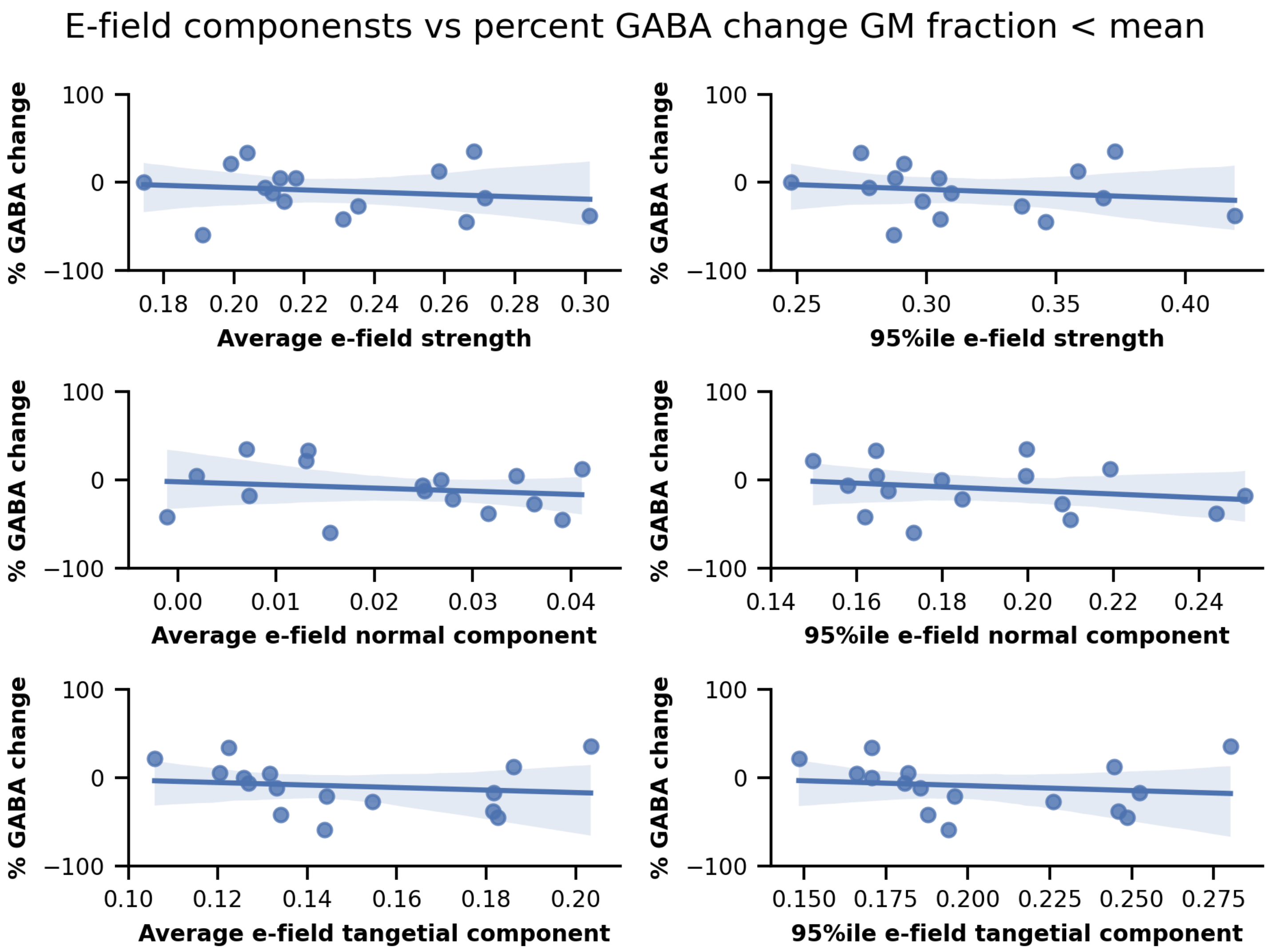


*Supplementary figure 6: Association between all the E-field estimates (magnitude, normal and tangent; mean and 95th percentile), in the temporal MRS voxel, and the percent GABA change, only for participants with grey matter partial volume estimates lower than the mean.*

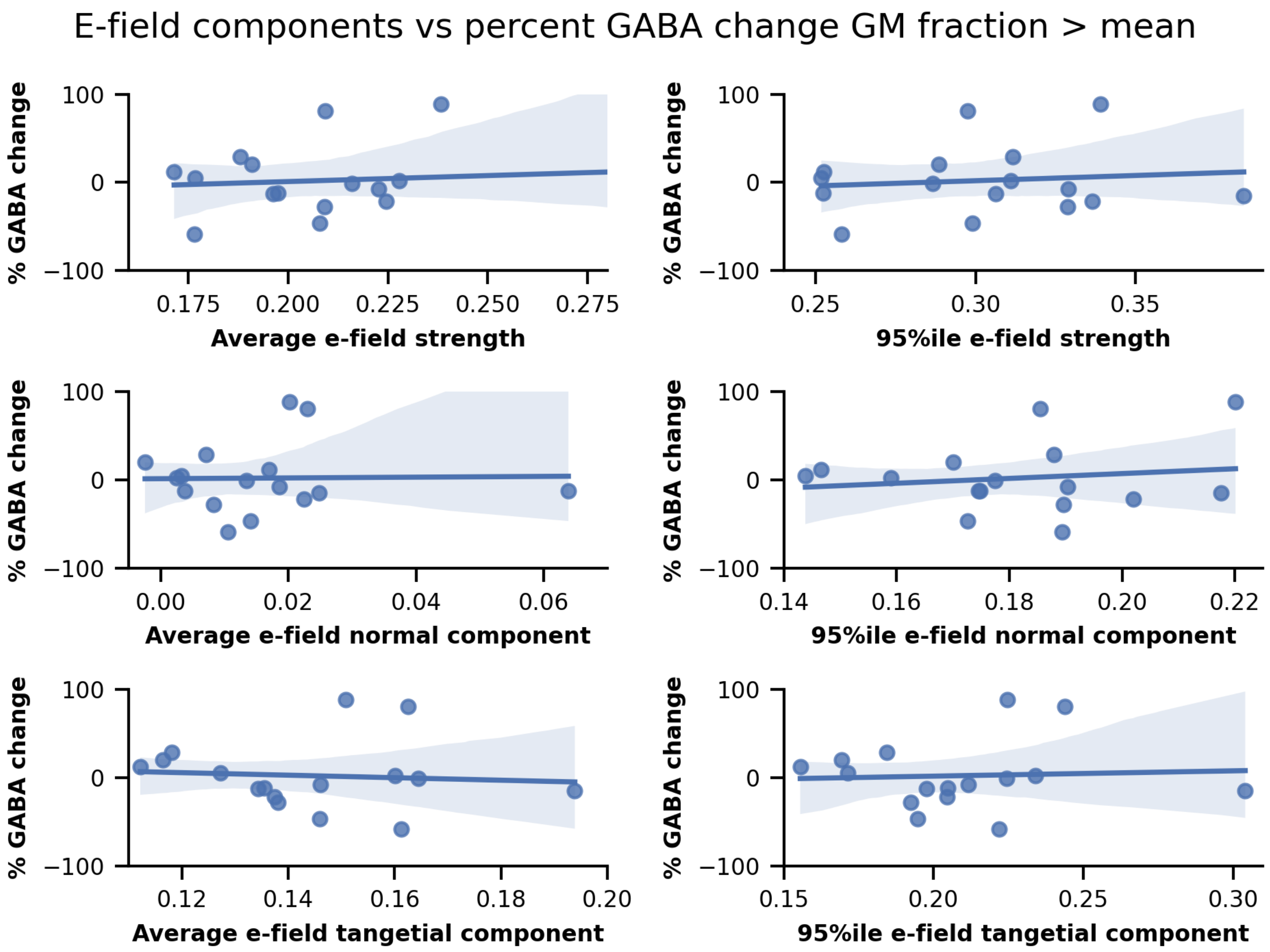


*Supplementary figure 7: Association between all the E-field estimates (magnitude, normal and tangent; mean and 95th percentile), in the temporal MRS voxel, and the percent GABA change, only for participants with grey matter partial volume estimates greater than the mean.*

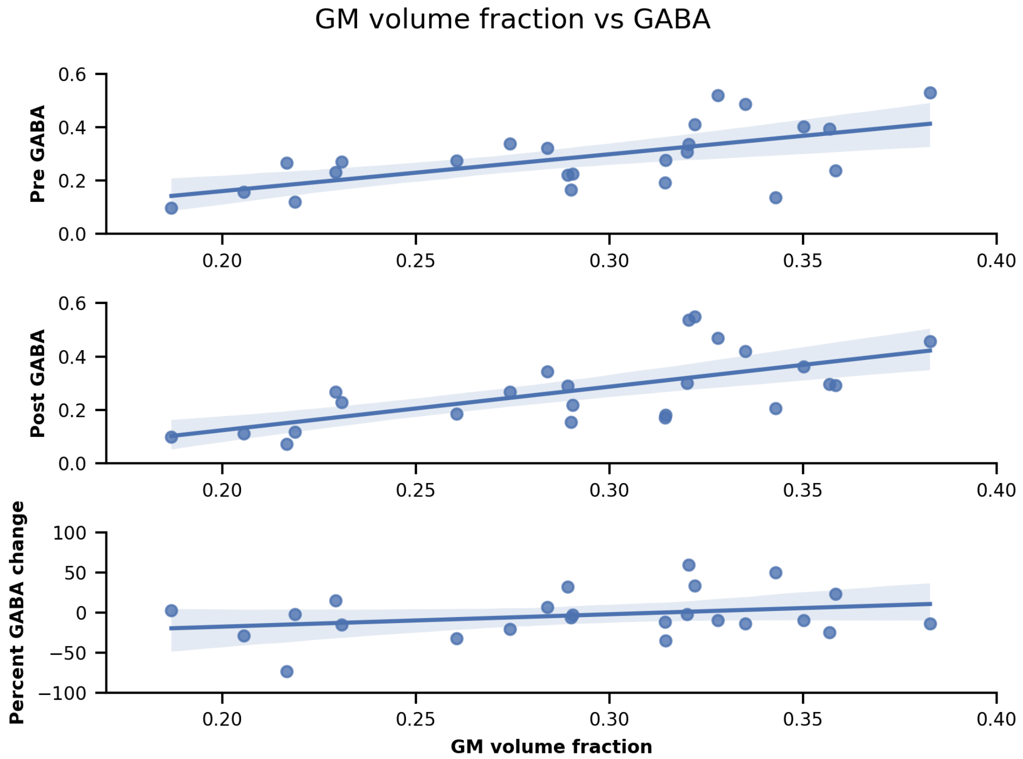


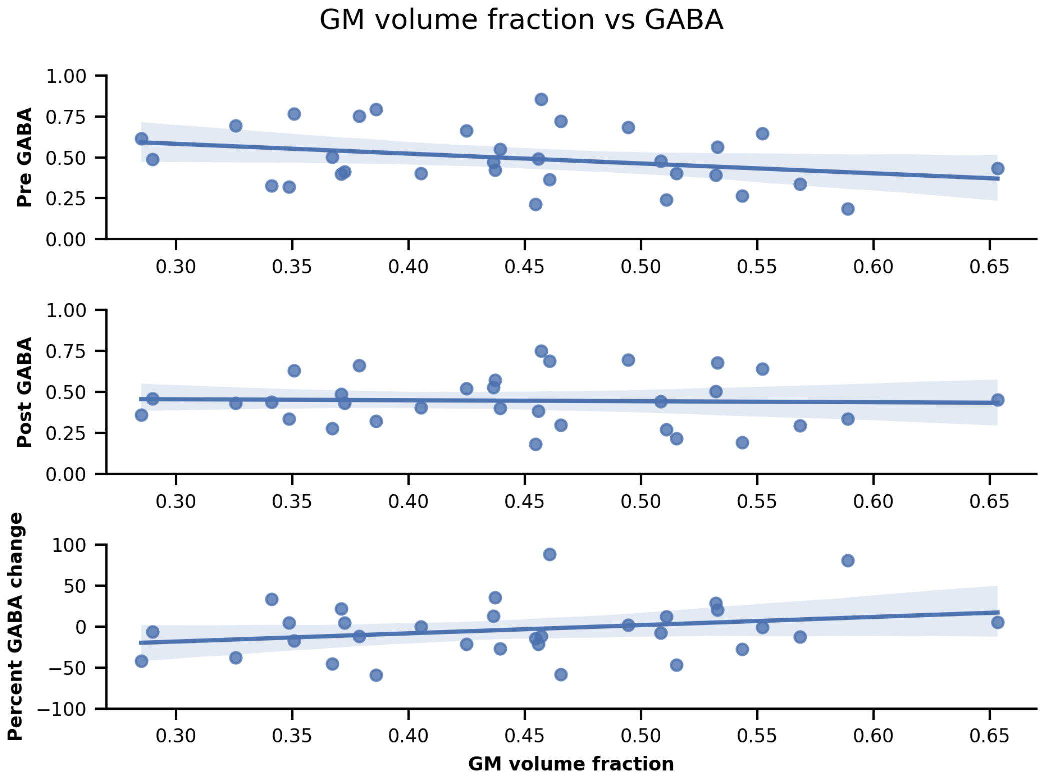


*Supplementary figure 8: Gaba change vs GM volume fraction in M1 (top) and temporal (bottom) area*

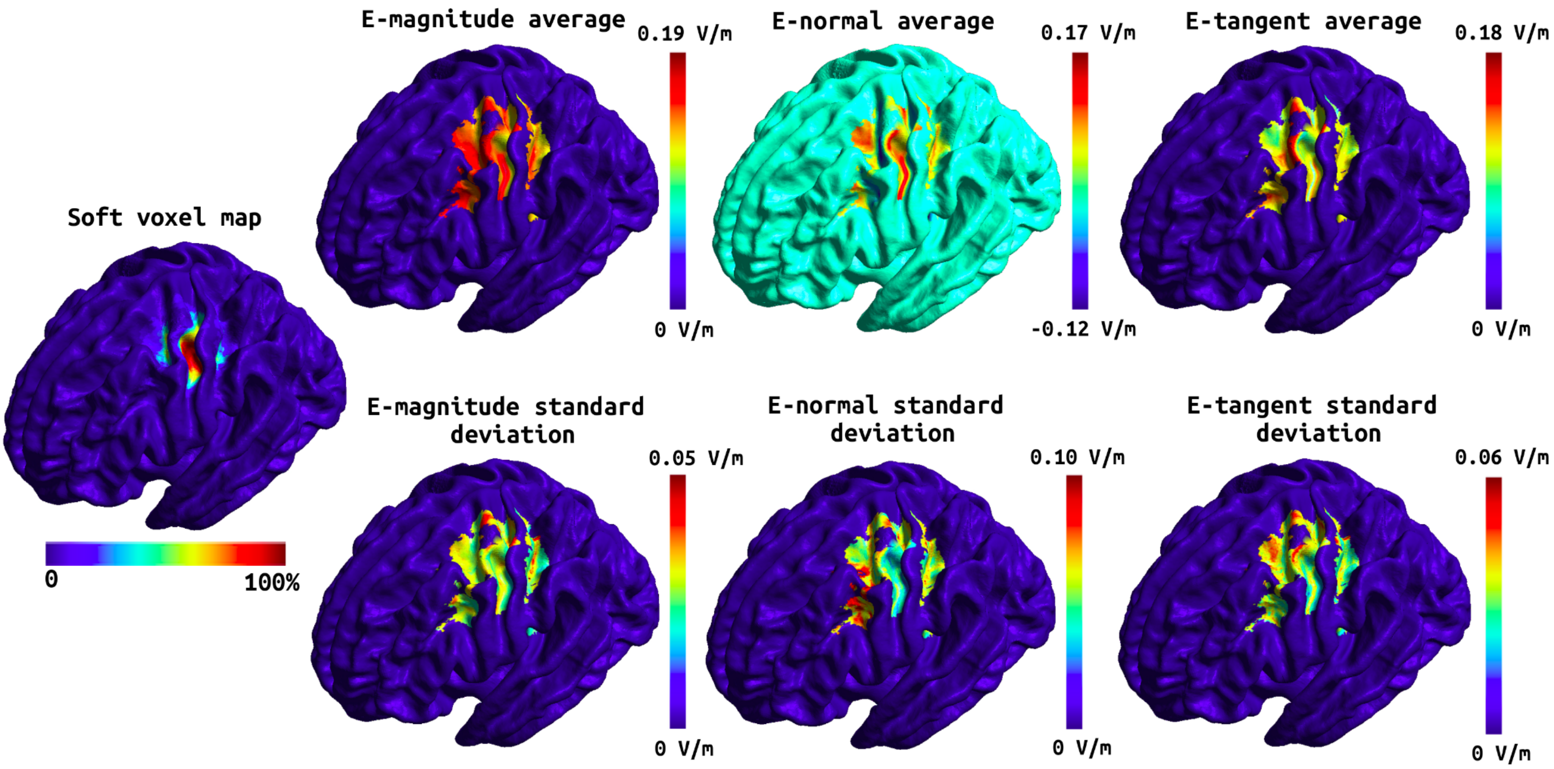


*Supplementary figure 9: Soft voxel map (leftmost) and the mean and standard deviation of the E-field components over the subjects in fsaverage space for the tDCS stimulation targeting the M1. First column: the mean E-field magnitude (top) and its standard deviation (bottom). Second column: the mean E-field normal component (top) and its standard deviation (bottom). Third column: the mean E-field tangential component (top) and its standard deviation (bottom).*

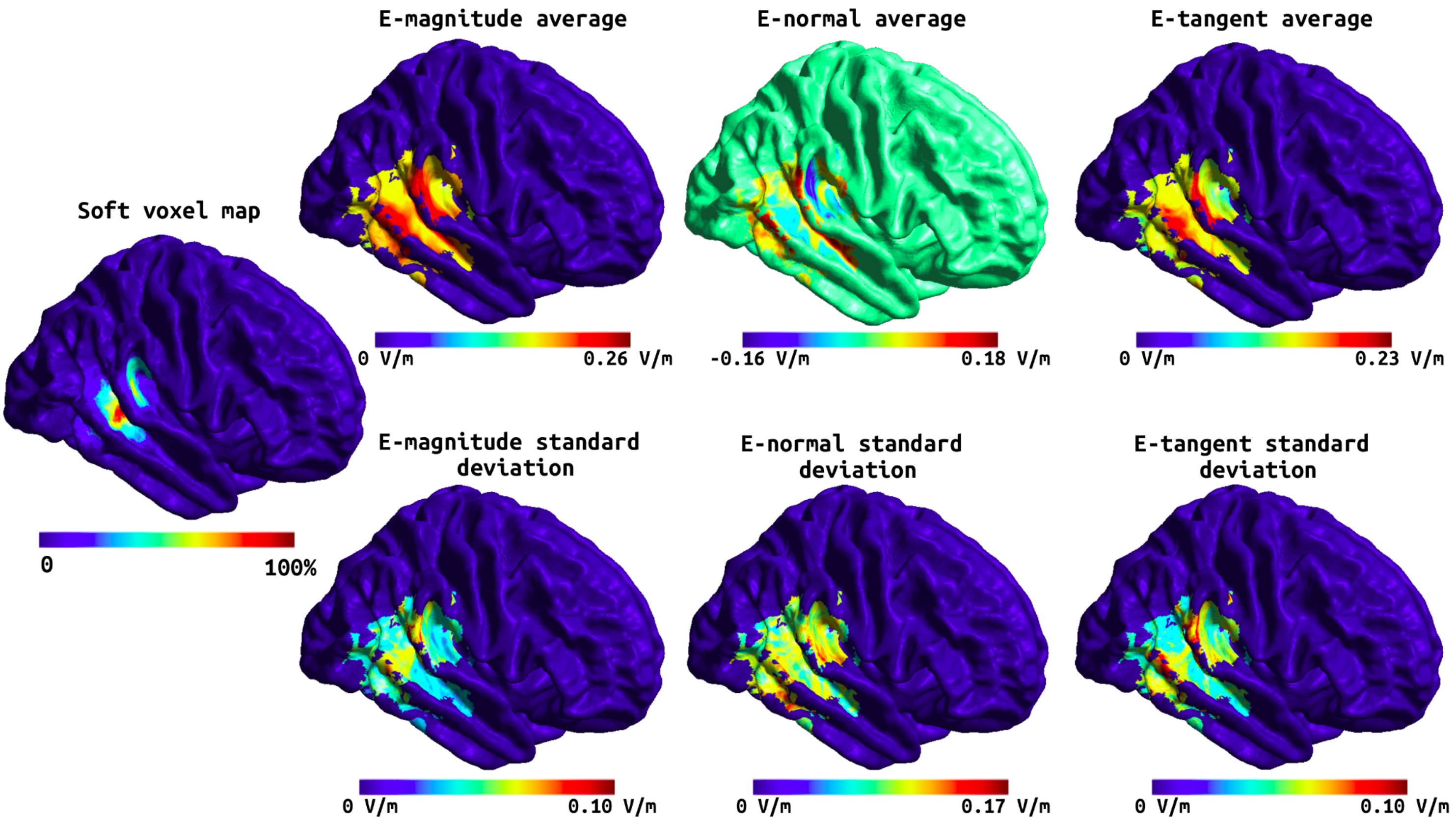


*Supplementary figure 10: Soft voxel map (leftmost) and the mean and standard deviation of the E-field components over the subjects in fsaverage space for the tDCS stimulation targeting the temporal cortex. First column: the mean E-field magnitude (top) and its standard deviation (bottom). Second column: the mean E-field normal component (top) and its standard deviation (bottom). Third column: the mean E-field tangential component (top) and its standard deviation (bottom).*

*Supplementary table 1: MRS data quality (included participants only)*

| ***Dataset*** | ***SNR*** | ***CRLB*** | ***NAA fwhm*** | ***Model fwhm*** | ***Creatine fwhm*** |
|---|---|---|---|---|---|
| ***Dataset 1 (unpublished)*** | *50.5 ± 5.8* | *21.3 ± 4.8* | | *9.6 ± 2.5* | |
| ***Koolschijn et al. 2019*** | *46.2 ± 9.6* | *18.6 ± 7.3* | | *10.9 ± 1.4* | |
| ***Bachtiar et al. 2018*** | *46.8 ± 6.8* | *20.1 ± 7.2* | | *9.9 ± 2.0* | |
| ***Barron et al. 2016*** | *51.2 ± 6.8* | *20.0 ± 6.4* | | *9.9 ± 1.3* | |
| ***Bachtiar et al. 2015*** | | | *9.1 ± 0.6* | | *8.3 ± 0.3* |

*SNR - signal to noise ratio, CRLB* - Cramér-Rao lower bounds, fwhm - full width at half maximum
SNR, CRLB and NAA fwhm were used for quality control during the analysis. We also provide the Model fwhm, estimated using LCModel for Semi-LASER data and Creatine fwhm, estimated using Gannet for MEGA-PRESS data since these are approximately comparable.

*Supplementary table 2: M1 model results*

| E-field variable | Main effects | | | Two-way interactions | | | Three-way interaction |
|---|---|---|---|---|---|---|---|
| | **Time** | **Efield** | **Grey matter volume** | **Time*Efield** | **Time*Grey matter volume** | ***Efield*Grey matter volume*** | ***Time*Efield*Grey matter volume*** |
| **Mean: magnitude** | t(24) = -3.35 **p=0.003*** | t(30.27) = -1.89 p = 0.07 | t(30.27) = -1.66 p = 0.107 | t(24) = 3.24 **p = 0.003*** | t(24) = 3.64 **p = 0.001*** | *t(30.27) = 2.16* ***p = 0.039**** | *t(24) = -3.55* ***p = 0.002**** |
| **95$^{th}$ percentile: magnitude** | t(24)= -3.31 **p=0.003*** | t(30.62) = -1.77 p = 0.086 | t(30.62) = -1.62 p = 0.115 | t(24) = 3.18 **p = 0.004*** | t(24) = 3.53 **p = 0.002*** | t(24) = 2.01 p = 0.052 | *t(24) = -3.41* ***p = 0.002**** |
| **Mean: normal component** | t(24) = -0.37 p = 0.713 | t(33.66) = -0.40 p= 0.69 | t(33.66) = 0.39 p= 0.698 | t(24) = 0.22 p = 0.824 | t(24) = 0.42 p = 0.677 | t(33.66) = 0.30 p = 0.767 | t(24) = -0.31 p = 0.763 |
| **95$^{th}$ percentile: normal component** | t(24) = -3.46 **p=0.002*** | t(29.56) = -1.40 p = 0.172 | t(29.56) = -1.14 p = 0.263 | t(24) = 3.38 **p = 0.002*** | t(24) = 3.73 **p = 0.001*** | t(29.56) = 1.53 p = 0.137 | *t(24) = -3.67* ***p = 0.001**** |
| **Mean: tangential component** | t(24) = -2.82 **p=0.010*** | t(33.05) = -2.34 **p = 0.026*** | t(33.05) = -2.12 **p = 0.042*** | t(24) = 2.66 **p = 0.014*** | t(24) = 2.96 **p = 0.007*** | t(33.05) = 2.72 **p = 0.010*** | t(24) = -2.82 **p = 0.009*** |
| **95$^{th}$ percentile: tangential component** | t(24) = -2.37 **p=0.026*** | t(32.48) = -1.70 p = 0.100 | t(32.48) = -1.53 p = 0.136 | t(24) = 2.17 **p = 0.041*** | t(24) = 2.66 **p = 0.014*** | t(32.48) = 2.13 **p = 0.041*** | t(24) = -2.47 **p = 0.021*** |

*Supplementary table 3: M1 posthoc tests for significant three-way interactions*

| Efield variable | $25^{th}$ percentile Grey matter volume | $75^{th}$ percentile Grey matter volume |
|---|---|---|
| **Mean: magnitude** | Chisq(1) = 1.78<br>p = 0.182 | Chisq(1) = 12.91<br>**p < 0.001*** |
| **$95^{th}$ percentile: magnitude** | Chisq(1) = 1.93<br>p = 0.165 | Chisq(1) = 9.72<br>**p = 0.002*** |
| **Mean: normal component** | - | - |
| **$95^{th}$ percentile: normal component** | Chisq(1) = 2.95<br>p = 0.086 | Chisq(1) = 14.69<br>**p < 0.001*** |
| **Mean: tangential component** | Chisq(1) = 2.00<br>p = 0.157 | Chisq(1) = 5.31<br>**p = 0.021*** |
| **$95^{th}$ percentile: tangential component** | Chisq(1) = 0.31<br>p = 0.581 | Chisq(1) = 7.57<br>**p = 0.006*** |

*Supplementary table 4: Temporal model results*

| **Efield variable** | **Main effects** | | | **Two-way interactions** | | | **Three-way interaction** |
|---|---|---|---|---|---|---|---|
| | **Time** | **Efield** | **Grey matter volume** | **Time*Efield** | **Time*Grey matter volume** | ***Efield*Grey matter volume*** | ***Time*Efield*Grey matter volume*** |
| **Mean: magnitude** | t(28) = 0.87<br>p= 0.393 | t(44.87) = 1.67<br>p = 0.102 | t(44.87) = 1.40<br>p = 0.170 | t(28) = -1.04<br>p = 0.310 | t(28) = -1.03<br>p = 0.314 | *t(44.87) = -1.53*<br>*p = 0.133* | *t(28) = 1.16*<br>*p = 0.255* |
| **95th percentile: magnitude** | t(28) = 1.54<br>p= 0.135 | t(43.64) = 2.06<br>**p = 0.045*** | t(43.64) = 1.82<br>p = 0.076 | t(28) = -1.71<br>p = 0.098 | t(28) = -1.79<br>p = 0.085 | *t(43.64) = -1.97*<br>*p = 0.055* | *t(28) = 1.92*<br>*p = 0.065* |
| **Mean: normal component** | t(28) = 0.54<br>p= 0.591 | t(44.90) = 0.36<br>p = 0.717 | t(44.90) = -0.59<br>p = 0.557 | t(28) = -1.22<br>p = 0.231 | t(28) = -0.47<br>p = 0.639 | *t(44.90) = -0.61*<br>*p = 0.523* | *t(28) = 1.08*<br>*p = 0.291* |
| **95th percentile: normal component** | t(28) = 1.34<br>p= 0.193 | t(44.17) = 1.70<br>p = 0.097 | t(44.17) = 1.34<br>p = 0.189 | t(28) = -1.48<br>p = 0.151 | t(28) = -1.32<br>p = 0.198 | *t(44.17) = -1.49*<br>*p = 0.144* | *t(28) = 1.43*<br>*p = 0.164* |
| **Mean: tangential component** | t(28) = 0.59<br>p= 0.564 | t(46.27) = 1.95<br>p = 0.057 | t(46.27) = 1.49<br>p = 0.144 | t(28) = -0.71<br>p = 0.483 | t(28) = -0.50<br>p = 0.618 | *t(46.27) = -1.69*<br>*p = 0.099* | *t(28) = 0.62*<br>*p = 0.542* |
| **95th percentile: tangential component** | t(28) = 0.68<br>p= 0.501 | t(46.02) = 2.15<br>**p = 0.037*** | t(46.02) = 1.77<br>p = 0.084 | t(28) = -0.82<br>p = 0.422 | t(28) = -0.64<br>p = 0.530 | *t(46.02) = -1.99*<br>*p = 0.053* | *t(28) = 0.76*<br>*p = 0.454* |
